# Population diversification in a yeast metabolic program promotes anticipation of environmental shifts

**DOI:** 10.1101/002907

**Authors:** Ophelia S. Venturelli, Ignacio Zuleta, Richard M. Murray, Hana El-Samad

## Abstract

Delineating the strategies by which cells contend with combinatorial changing environments is crucial for understanding cellular regulatory organization. When presented with two carbon sources, microorganisms first consume the carbon substrate that supports the highest growth rate (e.g. glucose) and then switch to the secondary carbon source (e.g. galactose), a paradigm known as the Monod model. Sequential sugar utilization has been attributed to transcriptional repression of the secondary metabolic pathway, followed by activation of this pathway upon depletion of the preferred carbon source. In this work, we challenge this notion. Although *Saccharomyces cerevisiae* cells consume glucose before galactose, we demonstrate that the galactose regulatory pathway is activated in a fraction of the cell population hours before glucose is fully consumed. This early activation reduces the time required for the population to transition between the two metabolic programs and provides a fitness advantage that might be crucial in competitive environments. Importantly, these findings define a new paradigm for the response of microbial populations to combinatorial carbon sources.

## INTRODUCTION

Microbial cells are continuously bombarded by diverse and changing combinatorial environmental stimuli. To survive and reproduce, a cell must accurately detect, assess, and selectively respond to these signals. Specifically, in competitive and unpredictable environments, cells need to constantly integrate information about the nature and quantities of nutritional substrates to scavenge maximum nutritional value[1]. Organisms that can balance the anticipation of future environmental shifts without sacrificing the rate of reproduction by excess metabolic burden exhibit a fitness advantage. However, optimal metabolic strategies for achieving this balance have not been thoroughly explored.

Studies of the response of microbial cells to the availability of multiple sugars has a long history, starting with the seminal work of Dienert in yeast[2,3] and Monod in bacteria[4,5]. When presented with both glucose and galactose, microbial cells consume these carbon substrates in a sequential manner rather than simultaneously metabolizing both, resulting in two separate growth phases[5]. In the first phase, cells preferentially metabolize the sugar on which they can grow the fastest (glucose in this case). Upon glucose depletion, cells transition to metabolizing the less preferred sugar (galactose). This response, classically known as “catabolite repression”, posits that the synthesis of the enzymes needed to metabolize the less preferred sugar is inhibited across the whole population. This inhibition is relieved by depletion of the preferred sugar, which triggers the diauxic shift. Crucially, in this model, the sequential consumption of the two sugars is generally attributed to the sequential expression of the enzymes needed for their metabolism[6]. In this work, we demonstrate that while *S. cerevisiae* cells indeed undergo sequential sugar consumption in the presence of combinations of glucose and galactose, the synthesis of the enzymes needed for the metabolism of galactose is not necessarily sequential. Specifically, we find that for a large combinatorial space of glucose-galactose inputs, a subpopulation of cells arises where the galactose transcriptional program is induced hours before the depletion of glucose. Intriguingly, these cells have a fully active galactose transcriptional program, yet they do not metabolize this sugar until glucose is exhausted. We demonstrate that this heterogeneous strategy is essential for rapid growth during the metabolic transition from glucose to galactose. These data suggest that the response of microorganisms to combinatorial environments may frequently involve diversification of phenotypes across a population. Furthermore, this strategy integrates direct environmental sensing with an anticipation of future environmental shifts. As such, it constitutes an elaboration on bet-hedging mechanisms that often rely on stochastic fluctuations to produce subpopulations with different phenotypes without a dominant input from the environment[7].

## RESULTS

**Figure 1.**
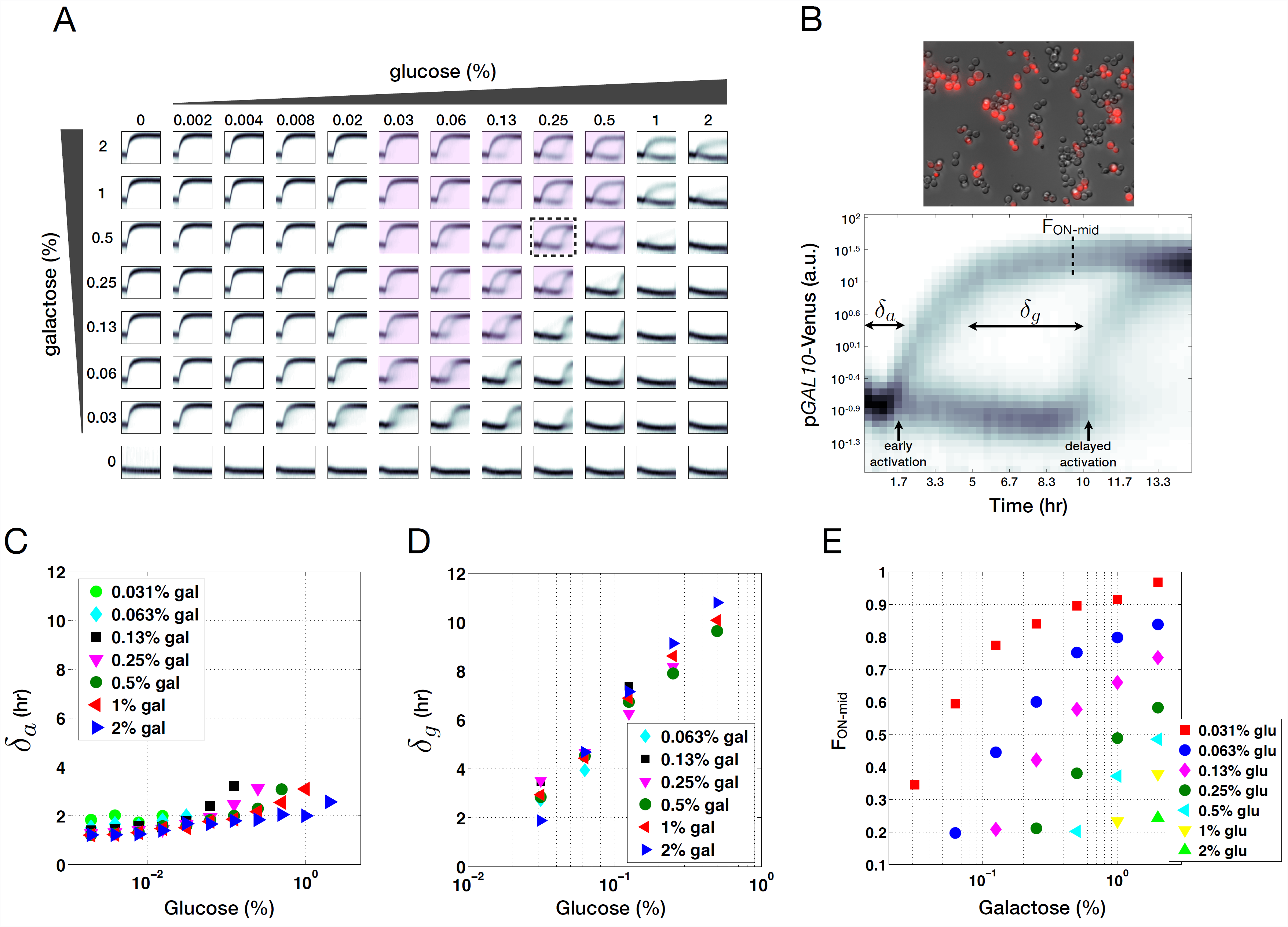
Dynamic responses of the GAL pathway to combinatorial inputs of glucose and galactose. **(A)** Single cell fluorescence distributions of pGAL10-Venus as a function of time in wild type *S. cerevisiae* obtained using automated flow cytometry for a wide range of glucose and galactose concentrations. In each subplot, the x-axis is time and the y-axis is fluorescence. Dashed box indicates the condition shown in panel **B** and highlighted conditions were used to quantify the duration of bimodality (δ_g_) in panel **D. (B)** Fluorescent microscopy image of pGAL10-Venus induced simultaneously with 0.25% glucose and 0.5% galactose at time zero and measured after 6 hours (top, corresponds to dashed box in panel **A**). pGAL10-Venus flow cytometry distributions as a function of time highlighting transient bimodality (bottom). δ_a_ represents the response time of the early activated subpopulation, δ_g_ represents the duration of bimodality for quantifiable conditions in panel **A** (highlighted conditions) and F_ON-mid_ denotes the fraction of cells in the ON state at the midpoint of the transient bimodal region (16). **(C)** Relationship between initial glucose levels and δ_a_ for a range of initial galactose concentrations. **(D)** Relationship between initial glucose concentrations and δ_g_ for different initial galactose levels. **(E)** Relationship between initial galactose levels and F_ON-mid_ for different initial concentrations of glucose.

We studied the time-resolved response of a population of yeast cells to combinatorial inputs of glucose and galactose using our automated flow cytometry setup that measures gene expression approximately every 20 min for 14 hours (Fig. 1A)[8]. This technology enabled us to measure the galactose (GAL) pathway activity dynamics in single-cells using the epimerase *GAL10* promoter (pGAL10) driving Venus (YFP) and to dissect the quantitative growth patterns of the microbial culture.

For sufficiently low glucose concentrations, pGAL10 induced as a single monomodal distribution. By contrast, pGAL10 did not activate over the course of the experiment for glucose concentrations significantly higher than those of galactose. These behaviors recapitulate previously observed phenotypes[9,10]. However, for a large spectrum of combinatorial glucose-galactose inputs aggregating around the regime of equal concentration of these two sugars, we observed the emergence of a bimodal gene expression response in which only a fraction of the population induced pGAL10 (Fig. 1A,B). In this regime, bimodality was transient since the cohort of OFF cells uniformly switched ON following a delay. The promoters of the galactokinase *GAL1* (pGAL1), permease transporter *GAL2* (pGAL2) and transferase *GAL7* (pGAL7) exhibited similar gene expression patterns, indicating that this transient bimodality was a general feature of the GAL pathway in response to a mixture of glucose and galactose (Supplementary Fig. 1).

**Figure 2.**
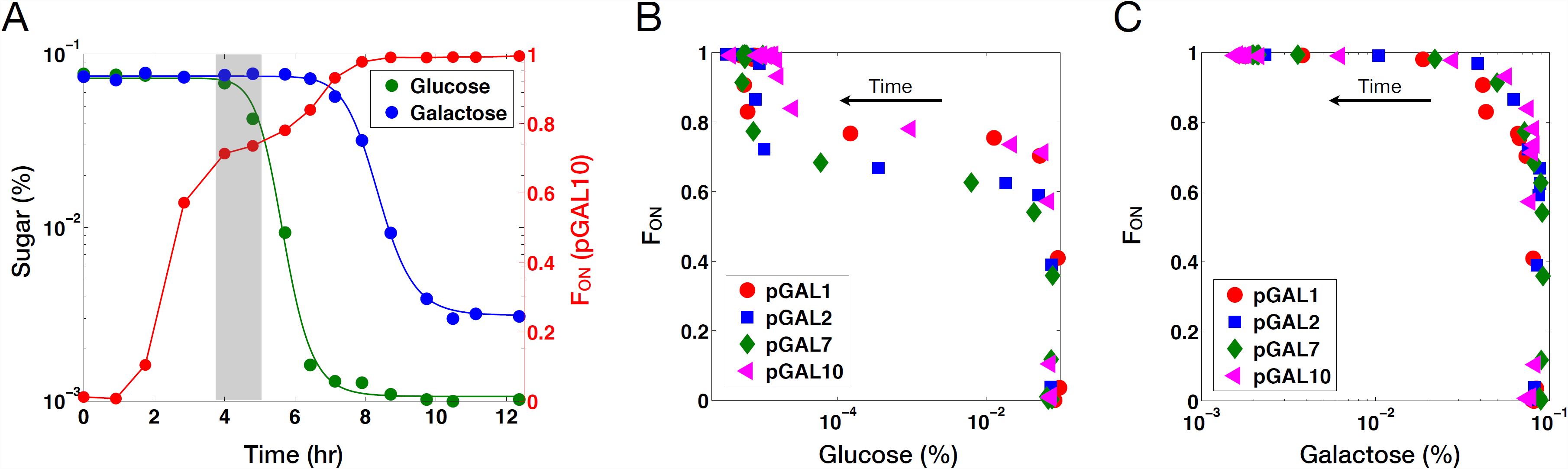
Glucose is consumed before galactose despite the presence of a subpopulation of cells with an active GAL pathway. Wild type cells were exposed to 0.1% glucose and 0.1% galactose simultaneously at time zero. **(A)** Representative dynamic measurements of glucose, galactose and the fraction of ON cells (F_ON_) for wild type expressing p*GAL10*-Venus. Highlighted box indicates plateaued region. Lines represent fitted Hill functions. **(B)** Scatter plot of glucose concentrations and F_ON_ for the *GAL1*, *GAL2*, *GAL7* and *GAL10* promoter fusions to Venus measured over time. **(C)** Scatter plot of galactose concentrations and F_ON_ for p*GAL1*, p*GAL2*, p*GAL7* and p*GAL10*.

Stochastic switching between the ON and OFF states in the transient bimodality region would generate a population of cells with intermediate fluorescence levels. This is due to the slow dynamics of protein synthesis and the high stability of fluorescent proteins, resulting in the decay rate of the fluorophore being dominated by dilution after cell division. However, in our data, the ON and OFF subpopulations were clearly separated from each other in the bimodal region and we did not detect cells of intermediate fluorescence values. In addition, for a given dual-sugar input, the fraction of GAL ON cells did not change significantly over time in the bimodality region (highlighted box in Supplementary Fig. 2A,B). Taken together with a previous study demonstrating that the GAL system can only exhibit stochastic transitions between states in the absence of the *GAL80* negative feedback loop but not in wild type[11], our data indicate that it is unlikely that cells are continuously switching between the ON and OFF states. Therefore, this phenomenon is distinguishable from previously observed stochastic switching between phenotypes[12,13].

**Figure 3.**
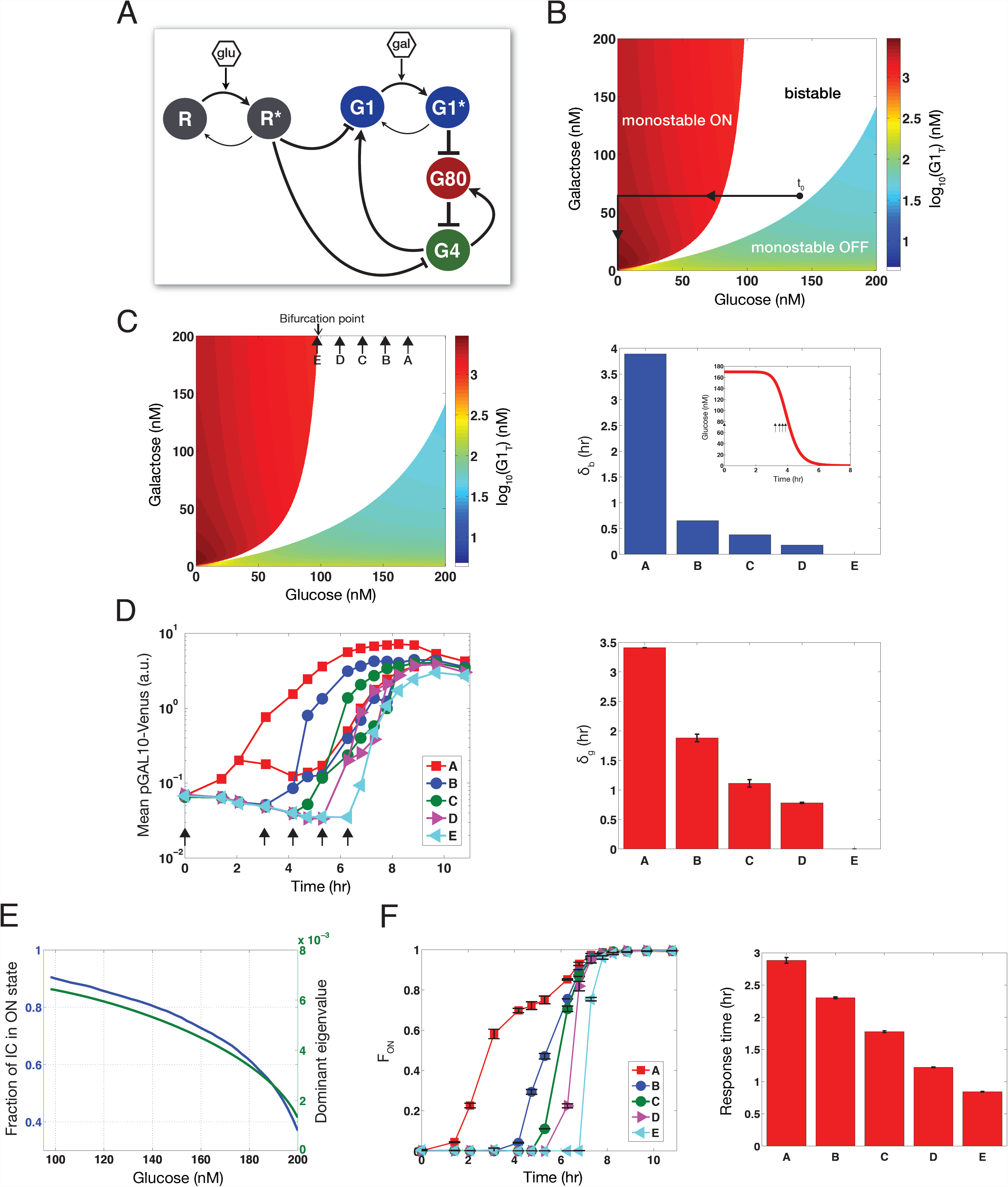
Computational model of GAL system with glucose and galactose inputs explains the origin of transient bimodality and predicts the dependence of the system on its inputs. **(A)** Schematic diagram of GAL circuit. Glu represents glucose and gal represents galactose. The activated molecules are represented by R* and G1*. Pointed and blunted arrows indicate activation and repression, respectively. **(B)** Bifurcation diagram at steady-state. Bistability is represented by white and colored regions denote monostability. Total concentration of G1 is denoted by G1_T_. Time zero is indicated by t_0_ and solid lines highlight a model trajectory of sequential consumption of the two sugars by a cell population. **(C)** Bifurcation diagram at steady-state (left). Arrows denote the addition of galactose at different times to a system that received glucose at t_0_. δ_b_ is computed as the time required for glucose to decay to the bifurcation point threshold (right). **(D)** Mean expression levels of ON and OFF subpopulations for the delayed galactose experiment (left) and duration of bimodality (δ_g_) (right). Arrows indicate the time of the galactose stimulus. **(E)** The fraction of initial conditions (IC) and the dominant eigenvalue for the ON equilibrium state as a function of glucose for 150 nM galactose. **(F)** Experimental measurements of F_ON_ in the galactose step experiment over time. Response time of F_ON_ for each condition (right). Error bars indicate one s.d. from the mean of two replicates.

Using a Gaussian mixture model (GMM) to deconvolve the two populations (see Methods), we quantified three measures of the response: the time to early activation for conditions with a detectable early activated population (δ_a_), the delay between early and late activation for conditions with transient bimodality (δ_g_, highlighted panels), and the fraction of ON cells quantified at the midpoint between the half-max of the early and delayed activation responses (F_ON-mid_) (Fig. 1B, see Methods). δ_a_ was modestly increased by glucose and reduced by galactose (Fig. 1C). By contrast, δ_g_, showed a substantial linear increase as a function of initial glucose (highlighted panels in Fig. 1A, Fig. 1D). However, δ_g_ was not significantly modified by the initial galactose concentration (Supplementary Fig. 3A). F_ON-mid_ significantly increased with the initial galactose level and was reduced by the initial glucose concentration for any given concentration of galactose (Fig. 1E). δ_g_ and F_ON-mid_ were modified in a set of mutants including regulators of the GAL pathway and glucose repression, suggesting that these phenotypes are modulated by a complex molecular program involving many factors (Supplementary Text, Supplementary Fig. 5A-D).

The existence of a subpopulation of cells in which the galactose transcriptional pathway was active in the transient bimodality regime suggested that the population might be consuming galactose concurrently with glucose. To test this hypothesis, we measured glucose, galactose and the fraction of ON cells (F_ON_) as a function of time in response to 0.1% glucose and 0.1% galactose, a condition in which the population exhibits a bimodal response. While these data recapitulated the known sequential order of sugar utilization, F_ON_ increased immediately following the dual-sugar stimulus and transiently plateaued before the cells consumed the available glucose (highlighted box in Fig. 2A and Supplementary Fig. 2). The initial concentration of glucose determined the duration of this plateau (Supplementary Fig. 2C). F_ON_ underwent a second increase to approximately 100% precisely at the time of total glucose depletion. Therefore, the timing of the delayed activation of the repressed subpopulation, and consequently the magnitude of δ_g_, seemed to be determined by the time of glucose depletion. In agreement with this hypothesis, δ_g_ was inversely related to the initial cell density N_0_, which modifies the rate of sugar consumption (Supplementary Fig. 3B). In addition, a population that received a first step of glucose and galactose, followed by an additional step input of glucose after 5 hours had a significantly larger δ_g_ than a population that received only the initial dual-sugar input, further corroborating the fact that δ_g_ is tuned by the concentration of glucose (Supplementary Fig. 3C). In contrast to δ_g_, F_ON-mid_ was approximately equal in conditions that received one or two steps of glucose, suggesting that the second glucose input did not induce substantial switching between OFF and ON states (Supplementary Fig. 3D).

**Figure 4.**
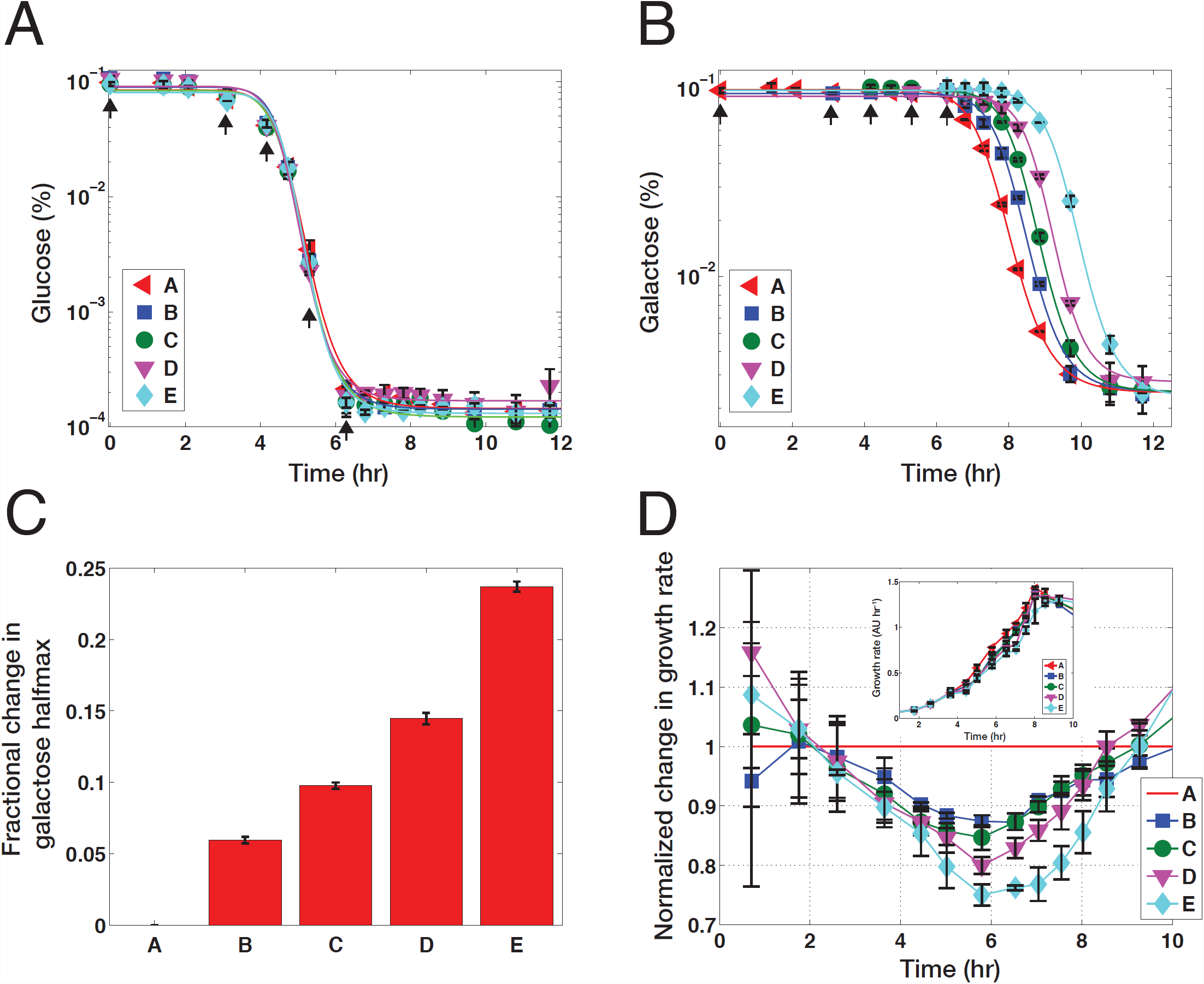
Early GAL pathway induction establishes a growth advantage during carbon source switch. Measurements of glucose, galactose and growth rates for delayed galactose experiment in which 0.1% galactose was added to a set of cultures at different times that had all received 0.1% glucose from time zero. Arrows indicate the time of the galactose stimulus. **(A)** Glucose concentrations as a function of time for each condition. **(B)** Galactose concentrations as a function of time for each condition. Lines represent fitted Hill functions. **(C)** Fractional change in the half-max of the galactose decay curves for each condition relative to condition A. **(D)** Normalized growth rates of conditions B-F compared to A (red line). Un-normalized growth rates for each condition (inset). Error bars represent one s.d. from the mean of two replicates.

Our results indicate that galactose consumption did not commence for hours despite the presence of a substantial subpopulation of GAL ON cells. Indeed, galactose consumption did not initiate until approximately 85% of cells were activated (Fig. 2C and Supplementary Fig. 4). Therefore, there is a broad regime where inhibition of galactose metabolism does not require transcriptional repression of the GAL genes. In this region, the population adopts a previously undocumented bimodal regulatory strategy of activated and repressed GAL states.

The absence of galactose metabolism does not result from decoupling galactose sensing and trafficking. In this inducer exclusion model[14], extracellular galactose is sensed, triggering GAL gene induction, but galactose does not permeate the cell due to glucose-dependent inhibition of galactose transport. However, Gal1p and Gal3p are the only known sensors of galactose and these proteins function intracellularly, indicating that a sufficient amount of galactose was entering the cells to induce pathway activation[15]. Furthermore, we observed a significant induction of pGAL2, accumulation of the fluorescently tagged Gal2 permease in the activated subpopulation, localization of Gal2p-Venus to the membrane in the presence of glucose and galactose and strong correlation between a Gal2 fluorescent protein fusion and pGAL10 in the presence of mixtures of glucose and galactose (Supplementary Figs. 1 and 6).

To further rule out the possibility of inducer exclusion as an explanation of our findings, we tested whether the level of Gal2p was limiting for the activation of the GAL pathway in the presence of glucose. To do so, we used a TET inducible promoter to vary the concentration of Gal2p in a strain deleted for the endogenous *GAL2* gene, and assessed the fraction of GAL ON cells (as quantified using a pGAL10-Venus reporter) in response to simultaneous addition of 0.5% galactose and a range of glucose levels (Supplementary Fig. 6D). We did not observe any dependence of the fraction of ON cells on aTc concentration, and hence on Gal2p levels, suggesting that inducer exclusion does not dominate pathway activation in the presence of glucose.

Taken together, our data suggest that in the cells where GAL genes are induced, transcriptional and metabolic control seem to be decoupled. These results counter a central premise of the Monod diauxic sugar model, predicated on the idea that expression of a secondary sugar pathway is repressed in the presence of a preferred carbon source, therefore blocking the utilization of the secondary sugar[4]. At the same time, our data are consistent with the observation that the inhibition of galactose consumption in response to a glucose pulse occurs on a timescale faster than can be explained by changes in transcriptional regulation or protein degradation, therefore making it unlikely that the catalytic degradation of Gal2p is the main effector of metabolic inhibition in the GAL pathway[16].

To understand how the structure of the GAL regulatory network could generate the observed transient bimodality in response to dual-sugar inputs, we constructed a simplified mathematical model of this circuit based on canonical knowledge about the galactose system (Supplementary Text)[17]. The GAL network has been shown to exhibit memory of galactose and glucose exposure, suggesting bistability as the source bimodality in this system[17,18]. In our model, the galactose input activates the signal transducer Gal1p (G1) forming G1*, which inhibits the repressor Gal80p (G80) from sequestering the transcriptional activator Gal4p (G4), thus leading to GAL gene activation (Fig. 3A). The inhibition of G80 liberates G4 to induce expression of G1 and G80, establishing a positive and negative feedback loop. Since glucose has been shown to reduce the activity of the GAL system, we coupled this model to an input of glucose[19]. We modeled GAL repression by glucose assuming that a repressor R (such as Mig1), can be activated by the glucose signal forming R*, which can then repress the promoters of *GAL1* and *GAL4*.

A salient qualitative feature of this mathematical model is that it can undergo a bifurcation from monostability to bistability as a function of its two inputs: glucose and galactose (Fig. 3B). In response to low glucose and high-galactose inputs, the model exhibits one steady-state, corresponding to the experimental ON state (high total G1 levels). For high glucose and low galactose inputs, the only steady-state corresponds to the OFF state (low total G1 levels). Similar concentrations of the two inputs produce two stable steady-states that correspond to the bimodality observed in the experiments.

This model also predicts the emergence and disappearance of bistability in the GAL system as a function of time for a given dual sugar input. By assuming that the system traverses a series of quasi-steady-states as a function of decaying sugar concentration (highlighted panels in Fig. 1), a given model trajectory crosses through a region of bistability, which is then transformed to monostability as glucose drops below a critical threshold (bifurcation point) due to cellular consumption (representative trajectory in Fig. 3B). This transition from bistable to monostable behavior at the glucose bifurcation point corresponds to the synchronized delayed activation of the repressed cohort of cells. The observed monomodal activation for sufficiently low glucose concentrations (left of highlighted panels in Fig. 1B) and significantly delayed activation (right of highlighted panels in Fig. 1B) are also explained by the model (Supplementary Fig. 7). Furthermore, the model indicates that if the glucose concentration were maintained above its value at the bifurcation point, for example by replenishment of glucose (Supplementary Fig. 3C), then the window of time where bistability exists in the system would be extended. This is precisely the case since cultures that received an initial pulse of glucose and galactose followed by a second pulse of glucose exhibited bimodality for a longer period of time compared to a culture that received only the initial sugar mixture (Supplementary Fig. 3C).

In addition to explaining the origin of transient bimodality, the model made predictions about different features of the system. First, the model had predictions about the role of feedback loops. Removing the *GAL80* feedback loop in the model augmented the range of glucose and galactose concentrations that produced bistability. This prediction was qualitatively consistent with our data showing that the range of glucose and galactose inputs that produced experimental bimodality was expanded in a strain lacking the Gal80p feedback loop (Supplementary Fig. 8).

The bifurcation hypothesis provided by the model also implied that the amount of time required for glucose to decrease to a threshold concentration corresponding to the bifurcation point in the system (δ_b_) should decrease if galactose is added at different times following the glucose input (rather than concomitantly with glucose—for example, 0, 3.2, 3.5 3.7 or 3.9 hours following an initial glucose step) (Fig. 3C). This increasing delay in galactose administration signifies a decreasing glucose concentration in the culture due to cellular consumption at the time of galactose addition, and hence a reduced window of time for bistability. The delayed activation response corresponds to the loss of bistability as glucose crosses a threshold bifurcation point. Hence δ_g_ and δ_b_ reflect similar properties of the system.

To experimentally test this prediction, we applied a step input of 0.1% galactose at different times to a set of cultures that had all received 0.1% glucose from time zero. In condition A, both sugars were added simultaneously at time zero, while in cultures B-E, galactose was added 3.1, 4.2, 5.3 and 6.3 hours following the glucose stimulus (arrows in Fig. 3D). Matching the trend of decreasing δ_b_ in the model (Fig. 3C), bimodality emerged at the time of the galactose input and δ_g_ contracted and eventually disappeared with the increased delay in this input (right panel in Fig. 3D, Supplementary Fig. 9).

Finally, the model indicated that the response time of the system to transition from the OFF to the ON state decreases as glucose decays. The system’s response time is dictated by both the domain of attraction and the magnitude of the dominant eigenvalue of the ON steady-state (Supplementary Text), which both increase as glucose decreases (Fig. 3E). Therefore, the model predicts that the response time of the fraction of ON cells should decrease in the delayed galactose experiment. Corroborating this insight, our experimental data demonstrated a decrease in the response time of F_ON_ with an increase in the delay of the galactose input (Fig. 3F,G). Therefore, in addition to providing a framework that explains the transition of the GAL system between different phenotypic modes, our model and its validated predictions demonstrate the intricate modulation of quantitative properties of this network by its environmental inputs. As we discuss below, this constitutes an important feature that distinguishes this strategy from previously documented stochastically dominated bet-hedging mechanisms.

We next probed the physiological impact of the observed anticipatory induction of the GAL regulatory program hours in advance of galactose consumption by analyzing the relationship between the timing of GAL pathway activation and the population’s growth rate and metabolism. To do so, we quantified the concentrations of glucose, galactose and growth rates for the different cultures that were subjected to delayed galactose inputs over time in the experiment described above. Irrespective of galactose timing, glucose decayed at a similar rate for all conditions (Fig. 4A). Despite the presence of glucose at the time of the galactose stimulus for conditions A-D, galactose consumption was delayed compared to the culture that received glucose and galactose simultaneously (Fig. 4B,C). Furthermore, the delay in galactose consumption was increased commensurately with the delay in galactose administration. Notably, during the metabolic shift between carbon sources, the population that received galactose simultaneously with glucose exhibited a transient growth rate advantage, reaching approximately 25% compared to the population that received this sugar after a 6.3-hour delay (E) (Fig. 4D, Supplementary Fig. 10). Since the growth rate is proportional to the current size of the population in exponential phase, the significance of this fitness difference increases with each cell generation. Overall, the delay in galactose input caused a monotonic increase in the transient growth defect, which was manifested as an increase in the “lag” time between the two phases of growth documented by Monod (Supplementary Fig. 10A)[4]. Importantly, the presence of galactose did not benefit the population of cells until total glucose depletion (Fig. 4D inset, Supplementary Figs. 10A and 11, Supplementary Text). Taken together, these data indicate that the induction of the GAL pathway many cell generations before these genes are required provides a transient fitness advantage during the shift between carbon sources.

This beneficial pre-emptive induction of the GAL pathway genes only occurs in a subset of the population. To investigate the tradeoffs that might motivate this bimodal induction, versus a uniform strategy in which all the cells in the population pre-emptively but coherently induce the GAL pathway, we sought to control GAL gene expression independently of galactose. To do so, we used an estradiol inducible Gal4 chimera in a strain lacking endogenous Gal4p[20,21]. In this strain, we could activate GAL gene expression on demand at specific times before glucose depletion in cultures subjected to 0.1% glucose and 0.1% galactose from time zero (Supplementary Fig. 12A).

Since the synthetic inducible system is not connected to the feedback structure of the natural circuit, GAL gene expression was monomodal (graded) as opposed to bimodal in this strain. In this case, early activation of the GAL pathway generated a lower consumption rate of glucose compared to late activation, demonstrating that constitutive GAL gene expression can inhibit glucose consumption (Supplementary Fig. 12B). The expression level of pGAL10 induced by this synthetic system was very similar to the expression level of pGAL10 in the wild type (Supplementary Fig. 13A). Therefore, the effects we observed are not likely to be a consequence of over or under expression of the GAL genes. Constitutive induction of the GAL pathway through over expression of Gal3p also reduced the glucose consumption rate (Supplementary Fig. 13, Supplementary Text). Together, these data highlight an important tradeoff that the system has to balance: induction of the GAL genes before they are required results in faster galactose consumption upon glucose depletion, facilitating the transition between carbon substrates. At the same time, wholesale induction of these genes across the entire population comes at the cost of a reduced rate of glucose consumption. In agreement with this observation, the repressed subpopulation had approximately 20% faster growth rate on average than that of the activated subpopulation in the transient bimodal region in the wild type (Supplementary Fig. 14, Supplementary Text). However, in this bimodal regime, the glucose consumption rate of the whole population was not saliently reduced by GAL gene expression in a subpopulation of cells (Fig. 4A). Therefore, the bimodal strategy seems to be efficiently balancing the tradeoffs imposed by the pre-emptive induction of the galactose transcriptional program.

## DISCUSSION

In this work, we demonstrate that a combinatorial input of glucose and galactose triggers diverse regulatory states across a population of cells. This transient bimodality establishes the co-existence of two subpopulation of cells—one that prepares hours in advance for a future shift in carbon metabolism and a second that defers pathway activation over many cell generations until these genes are required. The fraction of cells that occupy each state is tuned by the dual-sugar mixture, standing in contrast to canonical models in which the output of a pathway is proportionally matched to the level of its inputs in all cells of the population[22,23]. This mechanism also seems to be an elaboration on bet-hedging processes where stochastic fluctuations diversify the phenotypic states of a population in the absence of an environmental trigger[7,13,24], although in some instances, environmental cues have been implicated in biasing the distribution of phenotypic states[25,26]. There are some examples of biological networks that seem to have evolved the ability to activate multiple distinct signaling pathways simultaneously in response to a single environmental input as a consequence of temporal correlations between different environmental signals[27,28]. The strategy we describe is also unique in that the response of the GAL system integrates direct environmental sensing with pre-emption of a future metabolic shift.

Although our data do not pinpoint the exact mechanism by which glucose inhibits the metabolism of galactose in GAL ON cells, previous studies hint that this inhibition might be mediated by the dominant glucose kinase Hxk2p. Indeed, glucose and galactose are consumed simultaneously in cells lacking Hxk2p[29]. Furthermore, in *S. cerevisiae*, glucose was shown to block the maltose pathway (MAL) by a novel mechanism at the signaling level, which is also linked to Hxk2p[30] and distinct from inducer exclusion. It is therefore possible that the GAL and MAL pathways share similar post-translational inhibitory mechanisms by glucose.

Our data reveal that the induction of the GAL pathway in a fraction of the *S. cerevisiae* population before depletion of its preferred sugar (glucose) provides a kinetic advantage by shortening the lag phase before growth can resume on the secondary sugar. Evolutionary tuning of the duration of the lag phase has been shown to be a crucial variable for fitness of microbial populations in fluctuating environments[31]. Therefore, it is tempting to speculate that the advantage we characterize may be substantial for cells facing competition from other species for limited resources[32,33]. In agreement with this hypothesis, heterogeneity in the expression of the Lac operon in *E. coli* has recently been shown to modify the growth rates of single cells during the transition from glucose to lactose metabolism[34]. Furthermore, combinatorial carbon sources have been shown to trigger genetic mutations that produce phenotypic population diversification in *E. coli*[35]. Future studies that probe the broad adoption of similar strategies in other microorganisms, including *S. cerevisiae,* may yield insights into the precise evolutionary advantages of this response, and explore its use as a general paradigm for survival and reproduction in complex competitive environments.

## MATERIALS AND METHODS

### Growth conditions and flow cytometry

Cells were grown in yeast peptone media for approximately 12 hours and then diluted to an optical density (OD) of approximately 0.3 prior to induction with glucose and galactose. Single-cell fluorescence was measured on a LSRII analyzer (BD Biosciences). A blue (488 nm) laser was used to excite YFP and emission was detected using a 530/30 nm filter. 1000-20,000 cells were collected for each dynamic measurement.

### Automated flow cytometry measurements

A 500 μl culture volume was used in 96-well plate format for the automated flow cytometry measurements as described in ref. 9. For each time point, a 30 μl sample was removed from the culture for measurement on the cytometer and 30 μl of fresh media containing the appropriate 1X concentration of glucose and galactose was used as replacement to maintain a constant culture volume.

### Flask measurements

A 60 ml culture volume was used for the flask experiments in which the sugar concentrations were quantified. Less than 5% of the total volume was removed over the course of the experiment to quantify the single cell fluorescence, sugar concentrations and absorbance at 600 nm (OD). OD was measured on a Nanodrop 2000c spectrophotometer (Thermo Scientific).

### Quantitative analysis of gene expression dynamics

The ratio of YFP fluorescence to side scatter was used to quantify the total fluorescence per cell. Flow cytometry distributions were analyzed using a Gaussian mixture model algorithm (GMM, MATLAB) and each distribution was classified as either unimodal or bimodal as described in ref. 19. The delay time δ_a_ was computed as the time required to reach the half-max of the mean of the activated subpopulation. δ_g_ was defined as the difference between the time required to reach the half-max of the mean of the activated and repressed subpopulations. The fraction of ON cells (F_ON_) was computed as the fraction of the cell population higher than a fluorescence threshold (10^−0.2^ a.u.) that corresponds to approximately the lowest density of single-cell fluorescence between the OFF and ON expression states. F_ON-mid_ was quantified at the midpoint between the half-max of the activated (δ_a_) and repressed subpopulations. The response time was defined as the time to reach the half-max of F_ON_ (F_ON_ = 0.5). At each time point, individual cells were assigned to the OFF and ON states using the F_ON_ threshold on gene expression described above. The subpopulation growth rates were computed as the slope of a line fit to the log_2_ of the number of cells that accumulated in the OFF and ON states over time.

### Sugar measurements

Glucose and galactose were measured using the Amplex Red glucose oxidase and galactose oxidase kits (Molecular Probes, Life Technologies). A Tecan Safire plate reader (Tecan) was used to quantify the fluorescence. A standard of known concentration for each sugar was used to determine the quantitative relationship between the fluorescence and sugar concentration.

### Microscopy measurements

Cells were attached to glass-bottom wells in a 96-well plate (Matrical) functionalized with concanavalin-A (Sigma). Fluorescence images were taken at room temperature on a Nikon Ti-E equipped with a Perfect Focus System and a Coolsnap HQ2 CCD camera (Photometrics).

### Computational modeling

We used custom made code for mathematical modeling written in MATLAB (Mathworks) and Mathematica (Wolfram Research). Details about the model construction are provided in the Supplementary Text. The domain of attraction of the ON steady-state was defined as the fraction of initial conditions that were assimilated by the ON equilibrium point and was determined by randomly sampling 5000 initial conditions using the Latin Hypercube Method[17]. A minimum and maximum bound on the concentration of each species was used based on the parameters of the model. The dominant eigenvalue was defined as the eigenvalue of smallest absolute value of the linearization at the ON equilibrium point.

## Acknowledgements

### ACKNOWLEDGEMENTS

We would like to thank Adam Rosenthal and Michael Chevalier for helpful discussions and Carol Gross, Sandy Johnson, and Zev Gartner for critical reading of this manuscript. This research was supported by the Institute for Collaborative Biotechnologies through grant W911NF-09-0001 from the U.S. Army Research Office to R.M.M., the NIGMS system biology center (P50 GM081879) to H.E.S., and the David and Lucille Packard Foundation to H.E.S.

### AUTHOR CONTRIBUTIONS

All authors designed the experiments. O.V. and I.Z. carried out the experiments. O.V. performed the computational modeling and analyzed the data. All authors contributed to data interpretation and writing of the manuscript.

